# High quality single amplicon sequencing method for illumina platforms using ‘N’ (0-10) spacer primer pool without PhiX spik-in

**DOI:** 10.1101/2020.06.05.134197

**Authors:** Tejali Naik, Mohak Sharda, Awadhesh Pandit

## Abstract

Illumina sequencing platform requires base diversity in initial 11 cycles for efficient cluster identification and color matrix estimation. This limitation yields low quality data for amplicon libraries having homogeneous base composition. Spike-in of PhiX library ensures base diversity but overall reduces the number of sequencing reads for data analysis. To overcome such low diversity issues during amplicon sequencing on illumina platforms we developed high throughput single amplicon sequencing method by introducing ‘N’ spacers in target gene amplification primers that are pooled for simple handling. We evaluated the efficiency of ‘N’ spacer primers by targeting bacterial 16S V3-V4 region, demonstrating heterogonous base library construction. Addition of ‘N’ spacer causes sequencing frame shift at every base that leads to base diversity and produces heterogenous high quality reads within single amplicon library. We have written a python script “MetReTrim” to trim the heterogenous ‘N’ spacers from the pre-processed reads. This method terminates the need for PhiX spike-in and allows for multiplexing of multiple samples, greatly reducing the overall cost and yields improved sequence quality.

## Introduction

Amplicon sequencing is an important and widely used tool for inferring the presence of taxonomic groups in microbial communities, detecting genetic variation embedded in complex and genetic backgrounds, and is far more cost-effective than untargeted sequencing when large amounts of undesired genetic material is present (Lundberg et al., 2013; Callahan et al., 2019). Illumina HiSeq and MiSeq sequencing platforms are extensively used for performing paired-end sequencing to generate millions of reads for amplified fragments of the 16S rRNA gene, the internal transcribed spacer (ITS) region and different marker genes (Holm et al., 2019). Illumina’s sequencing-by-synthesis technology uses fluorescently labelled reversible terminator-bound dNTPs. Red laser illuminates A and C and green laser illuminates G and T fluorophores. Different filters are employed to image and identify the four different nucleotides. The similar emission spectra of the fluorophores (A and C as well as G and T) and limitations of the filters increases chances of low base call quality and mismatch rate in homogeneous sequence libraries (Schirmer et al., 2015). Therefore, for effective template generation and accurate base-calling on Illumina platforms it is required to have nucleotide diversity (equal proportions of A, C, G, and T nucleotides) at each base position in a sequencing library (Muinck et al., 2017; illumina 2014). Libraries of low sequence diversity like 16s rRNA gene are highly homogenous and commonly spiked with high-diversity library such as PhiX, to alleviate the problem of homogenous signals generated across the entire flow cell. However, it reduces the overall sequence read throughput and multiplexing options because of it being a non-target (PhiX) library (Holm et al., 2019; Muinck et al., 2017). The base diversity in first few cycles, particularly in the first 11 bases of the amplicon, are crucial for identification of the sequencing clusters on the flow cell and color matrix estimation (Jensen et al., 2019; Holm et al., 2019). Even though the research field has progressed in successful sequencing of 16S rRNA with Illumina V3–V4 region primers, the problems of drop in read quality and inherent error rate still remain unresolved (Jensen et al., 2019). Another approach to deal with this issue is by sequencing libraries tagged with heterogeneity spacers at the 5’ end of the target gene amplicon during library preparation. The heterogeneity spacers are short sequences linked to index adaptors or to the gene specific amplification primers in the form of 0-7 bases and minimizes the need for PhiX spike-in to 10 % by introducing base complexity at the start of sequencing reads yielding high quality sequencing and increased multiplexing capacity (Muinck et al., 2017; Holm et al., 2019; Herbold et al., 2015; Kozich et al., 2013). However, designing primers or index adaptors consisting varying length of heterogeneity spacer with unique sequences for different types of amplicon libraries is a complex process due to the fact that every base sequenced at given time should contribute to diversity (A ∼25%, T ∼25%, G ∼25%, C ∼25%) during the sequencing run and also requires PhiX spike-in. The PhiX spike-in hinders the use of Miseq and Hiseq platforms to great levels. Another drawback is handling more number of heterogeneity primer pairs instead of a single gene specific primer pair. Amplification of target gene with 0-7, 1-6, 2-5 and so on combinations of primer pairs makes the experimental setup tedious and requires minimum 8 samples to be pooled for confirming base complexity.

To resolve the technical limitations of single amplicon sequencing on Illumina platforms and challenges encountered during heterogeneity spacer primer designing, we added ‘N’ nucleotides to the 5’ end of the gene specific primers for amplifying the gene. The ‘N’ nucleotide bases are added in 0-10 fashion in forward and reverse gene specific primers. Pool of ‘N’ (0-10) spacers-linked gene specific primers are used for amplification and library synthesis incorporating diversity within single library. In addition, the pool design reduces the number of primer combinations to single set compared to previous studies (Liyou et al., 2015; Jensen et al., 2019). This contributes to increased base diversity at each sequencing cycle in all the libraries that are multiplexed during a sequencing run on Illumina platform. Our ‘N’ (0-10) spacer primers form libraries with improved base diversity leading to a higher quality data. Since our method is devoid of PhiX spike-ins, this allows for sequencing of more number of samples and a reduction in the overall cost. Further, this strategy of using ‘N’ (0-10) spacer design can be simply adopted for generating high quality single locus amplicon sequencing in a high throughput manner on any illumina platform.

## Methods

### 1. Reagents

| Product | Supplier | Catalog Number |
| --- | --- | --- |
| Nuclease-free water | Ambion | AM993 |
| Ethanol | Merck kGaA | 64-17-5 |
| AMPure-XP Magnetic Beads | Beckman Coulter | A63881 |
| Ultrapure 1 MTris-HCL pH 8.0 | Invitrogen | 15568-025 |
| DNA High Sensitivity Qubit Reagents | Invitrogen | Q32854 |
| 2x KAPA HiFiHotStart Ready Mix | KapaBiosystems | KK2602 |
| Agilent 2100 DNA 1000 Bioanalyzer Kit | Agilent | 5067-1504 |
| Nextera XT Index kit V2 | Illumina | 15032350 |
| NaOH | Sigma-Aldrich | 1091361000 |
| MiSeq V3 reagent cartridge and Flow cell | Illumina | MS-102-3003 |
| pH Test Strips (7.0-14.0) | Sigma | P-4411 |
| 16S Amplicon PCR Forward ‘N’ spacer Primers | Sigma | Sigma |
| 16S Amplicon PCR Forward ‘N’ spacer Primers | Sigma | Sigma |

#### 1.1. Buffer

1.2.1 1mM Tris-HCL: In falcon tube add 500 µl of 1 M Tris-HCL pH 8.0 to 49.5 mL Nuclease free water. Prepare fresh and keep at RT.
1.2.2 0.1N NaOH: Add 50ul of 2N NaOH to 950ul of Nuclease free water. Check the pH using pH test strips. The pH should be >12.5.0.1 N NaOH should be freshly prepared for denaturing libraries.

### 2. Equipment

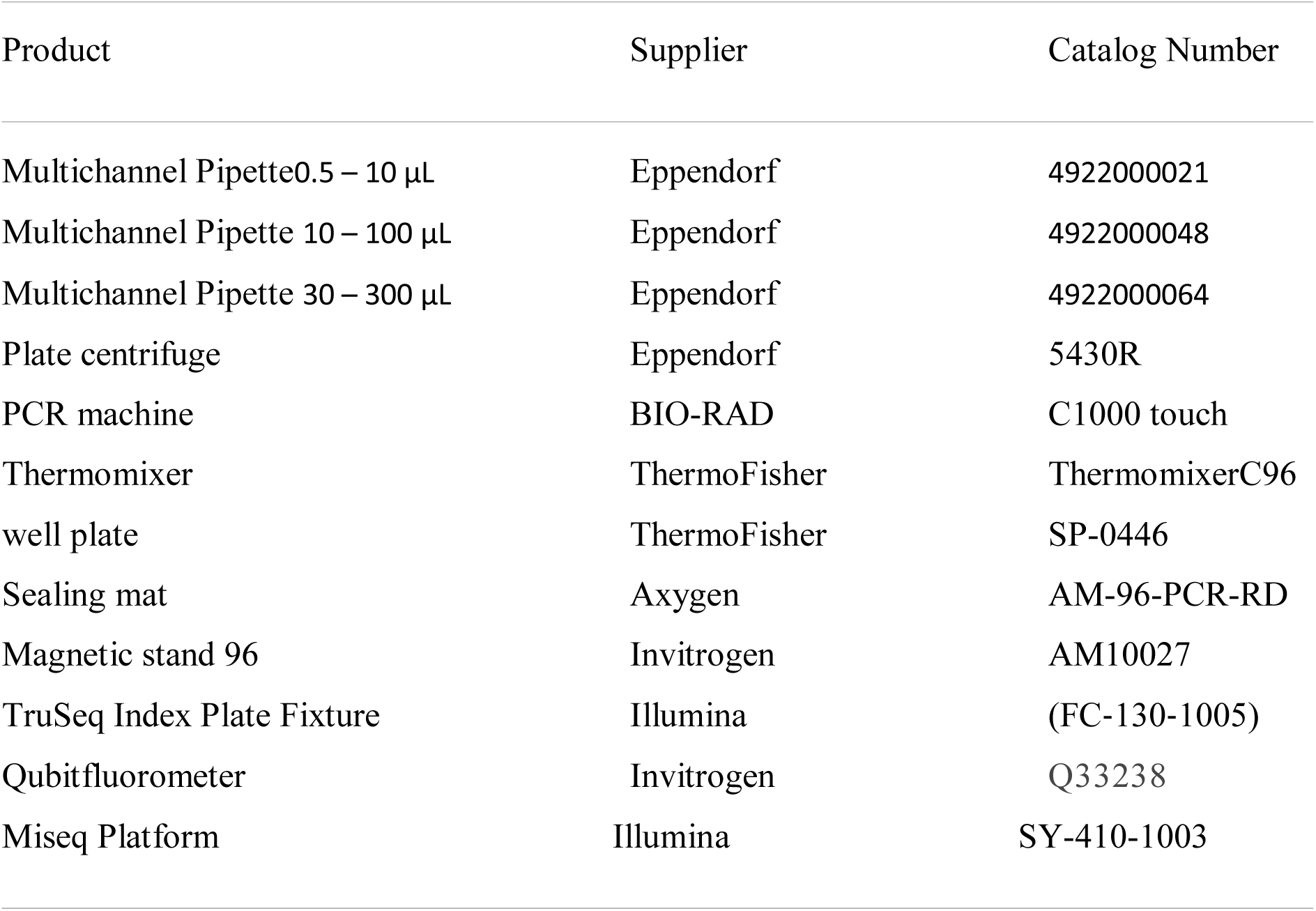

### 3. Experimental design

#### 3.1. Primer Design

Bacterial 16S V3-V4 region was targeted to study the efficiency of ‘N’ spacer primer. The primer contains Illumina adapter overhang sequence (blue), ‘N’ spacer region (red) and Target Gene specific primer (green). (Fig.1, Table 1 and Table 2). The primers were ordered as standard desalted PCR primers.

**Table 1:**
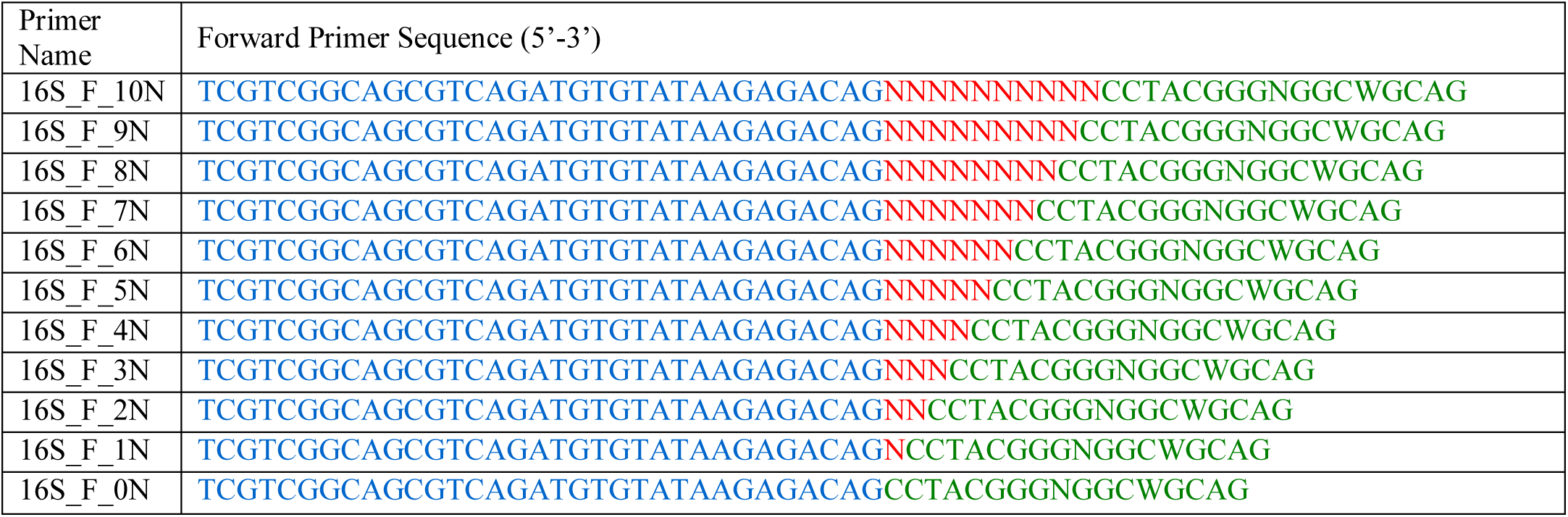
Forward ‘N’ spacer primers required for the First round PCR

**Table 2:**
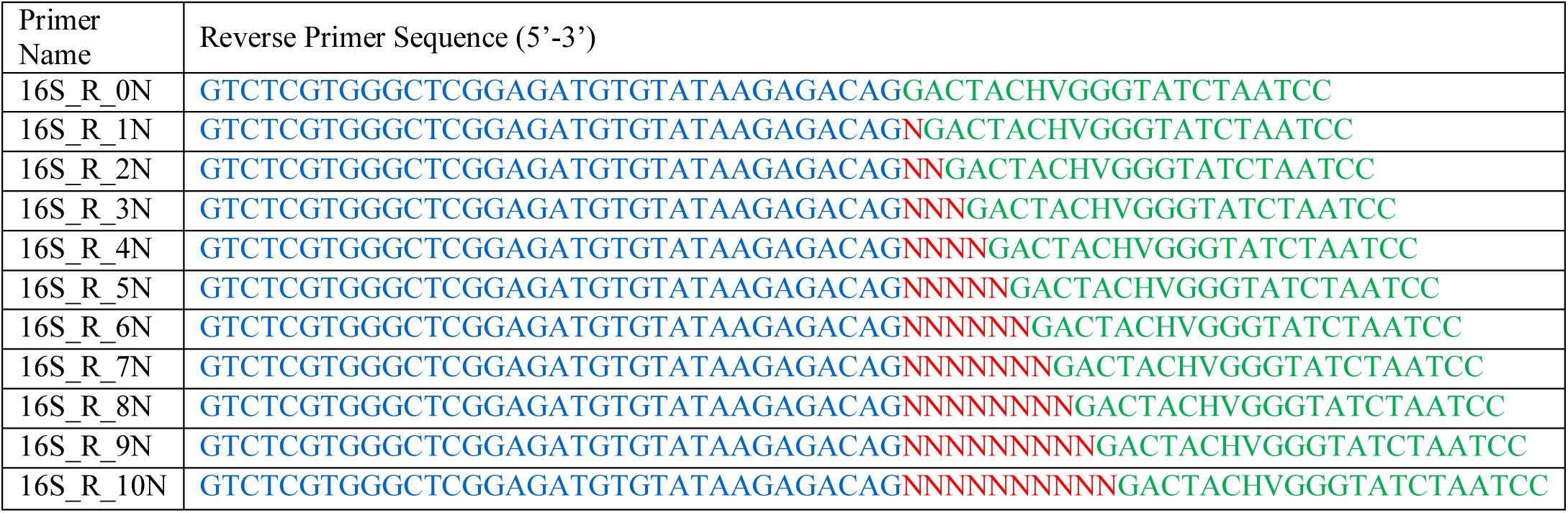
Reverse ‘N’ spacer primers required for the First round PCR

Illumina adapter overhang Sequence:

Forward overhang: 5’ TCGTCGGCAGCGTCAGATGTGTATAAGAGACAG Reverse overhang: 5’ GTCTCGTGGGCTCGGAGATGTGTATAAGAGACAG

Forward and Reverse primer stocks were diluted to 5uM and equal volumes of each forward and reverse primer were pooled together.

Critical Step: For freshly ordered primers it is recommended to check efficiency of each primer before pooling them. Perform PCR using control template and any combination of forward and reverse primer in total 11 PCR reaction setup.

#### 3.2. Starting Material

Genomic DNA 12.5 ng

### 4. Detailed Protocol

#### 4.1. First round PCR-Amplifying target gene

Assemble the following components per reaction

| Reagents | Per well |
| --- | --- |
| Microbial Genomic DNA (12.5 ng) | 2.5µL |
| Forward Primer Pool (5uM) | 1µL |
| Reverse Primer Pool (5uM) | 1µL |
| 2x KAPA HiFi HotStart ReadyMix | 12.5µL |
| Nuclease Free Water | 8µL |

Use 96 well plate for more samples.

Carry out the following PCR program:

i. 95°C for 3 min,
ii. 95°C for 30s, 55°C for 30s, 72°C for 30s for 25 cycles
iii. 72°C for 5 min, Hold at 10°C

[Optional] Check for PCR amplification by estimating the concentration of few amplicons on Qubit using DNA High Sensitivity Qubit Reagent.

#### 4.2. Amplicon Cleanup

4.2.1. Centrifuge the Amplicon PCR plate at 1,000 × g at 20°C for 1 minute.
4.2.2. Vortex the AMPure XP beads for 30 seconds to make sure that the beads are evenly distributed
4.2.3. Add 0.8X (20 µl) of AMPure XP beads to each well using multichannel pipette, mix the beads by gently pipetting entire volume 10-15 times. Seal plate and shake at 1800 rpm for 2 minutes.
4.2.4. Incubate the plate for 5 minutes at RT
4.2.5. Keep the plate on Magnetic stand for 5 minutes or until the supernatant is clear.
4.2.6. Without disturbing the plate kept on Magnetic stand, discard the supernatant using Multichannel Pipette.
4.2.7. With the Amplicon PCR plate on the magnetic stand, wash the beads by adding 150ul of freshly prepared 80% ethanol to each sample well. Incubate for 30 seconds. Carefully remove and discard the supernatant.
4.2.8. Repeat the above step for total two washes. Using multichannel pipette carefully remove the leftover ethanol from each well.
4.2.9. With the Amplicon PCR plate on the magnetic stand, air-dry the beads for 2 mins. Do not over dry the beads
4.2.10. Remove the Amplicon PCR plate from magnetic stand, using multichannel pipette resuspend the beads in 25 µl of 10 mMTris pH 8.0 in each well.
4.2.11. Mix the beads by gently pipetting entire volume 10-15 times to ensure proper resuspension of beads. Seal plate and shake at 1800 rpm for 2 minutes.
4.2.12. ncubate for 2 minutes at RT
4.2.13. Place the plate on the magnetic stand for 5 minutes or until the supernatant is clear.
4.2.14. Using a multichannel pipette, carefully transfer 22.5 µl of the supernatant to new 96 well PCR plate.
4.2.15. Quantify the amplicons using Qubit DNA HS reagent kit.

[Optional] Verify the amplicon size on Bioanalyzer using DNA 1000 chip. For V3-V4 region expected size after PCR is ∼550-560 bp.

#### 4.3. Second round PCR-Nextera XT Index Barcoding

This step adds Index 1 (i7) and Index 2 (i5) sequences to generate uniquely tagged libraries by amplifying the target gene amplicons using illumina Nextera XT Index Kit V2. The forward primer contains the P5 adapter end that binds to the flow cell, unique 8 nt index and the Illumina read 1 primer binding site. The reverse primers consist of the P7 adapter end that bind to the flow cell,unique 8 nt index and the Illumina read 2 primer binding site. Refer to Illumina’s Index Adapters Pooling Guide for selection of compatible primer combinations.

In TruSeq Index Plate Fixture arrange the Nextera XT Index 1 primer (i7) tubes horizontally from 1-12 fashion and Nextera XT Index 2 primer (i5) tubes vertically in 1-8 fashion. Transfer 1ul of purified product to a new 96 well PCR plate and place it on TruSeq Index Plate Fixture.

Set up the following reaction using Multichannel pipette for adding Index primer 1 and primer2:

| Reagents | Per well |
| --- | --- |
| Amplified Product | 1 μl |
| Nextera XT Index Primer 1 | 5 μl |
| Nextera XT Index Primer 2 | 5 μl |
| 2x KAPA HiFi HotStart ReadyMix | 25 μl |
| Nuclease Free water | 14 μl |

Gently pipette up and down 10-15 times and seal the plate with septa. Centrifuge the plate at 1,000 × g at 20°C for 1 minute.

Carry out the following PCR program:

i. 95°C for 3 min,
ii. 8 cycles of 95°C for 30s, 55°C for 30s, 72°C for 30s
iii. 72°C for 5 min, Hold at 6°C

#### 4.4. Index Amplicon Cleanup

4.4.1. Centrifuge the Amplicon PCR plate at 1,000 × g at 20°C for 1 minute.
4.4.2. Vortex the AMPure XP beads for 30 seconds to make sure that the beads are evenly distributed
4.4.3. Add 1X (50 µl) of AMPure XP beads to each well using multichannel pipette, mix the beads by gently pipetting entire volume 10-15 times. Seal plate and shake at 1800 rpm for 2 minutes.
4.4.4. Incubate the plate for 5 minutes at RT
4.4.5. Keep the plate on Magnetic stand for 5 minutes or until the supernatant is clear.
4.4.6. Without disturbing the plate kept on Magnetic stand, discard the supernatant using Multichannel Pipette.
4.4.7. With the Amplicon PCR plate on the magnetic stand, wash the beads by adding 150ul of freshly prepared 80% ethanol to each sample well. Incubate for 30 seconds. Carefully remove and discard the supernatant.
4.4.8. Repeat the above step for total two washes. Using multichannel pipette carefully remove the leftover ethanol from each well.
4.4.9. With the Amplicon PCR plate on the magnetic stand, air-dry the beads for 2 mins. Do not over dry the beads.
4.4.10. Remove the Amplicon PCR plate from magnetic stand, using multichannel pipette resuspend the beads in 37.5 µl of 10 mMTris pH 8.0 in each well.
4.4.11. Mix the beads by gently pipetting entire volume 10-15 times to ensure proper resuspension of beads. Seal plate and shake at 1800 rpm for 2 minutes.
4.4.12. Incubate for 2 minutes at RT
4.4.13. Place the plate on the magnetic stand for 5 minutes or until the supernatant is clear.
4.4.14. Using a multichannel pipette, carefully transfer 35 µl of the supernatant to new 96 well PCR plate.
4.4.15. **Quantify the libraries using Qubit DNA HS reagent kit**.
4.4.16. **Check the final library on a Bioanalyzer DNA 1000 chip to verify the size. Expected size for V3-V4 Region is ∼630-640bp**.
4.4.17. **Library Normalization and Pooling**

Calculate DNA concentration in nM, based on the size and concentration of library

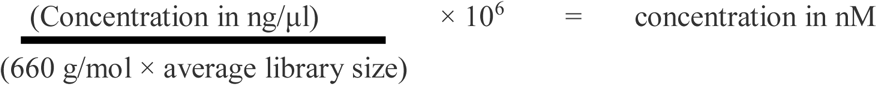

Dilute the final library using 10 mMTris pH 8.0 to 4 nM. Aliquot 5 μl from each diluted library and mix aliquots for pooling libraries with unique indices. Depending on coverage needs, 96 or more than 96 libraries can be pooled for one MiSeq run. Check the concentration of pooled library and calculate the nM considering the average size of all libraries. It should be ∼4nM

### 5. Library Denaturing and MiSeq Loading

5.1.1. Keep the Miseq Reagent for thawing at RT
5.1.2. Freshly prepare 0.2N NaOH
5.1.3. Keep the thawed HT1 buffer in Ice
5.1.4. Denature the Library

5.2. Combine the following in a 1.5 ml tube
  4 nM pooled library (5 μl)
  0.2 N NaOH (5 μl)
5.3. Mix with pipette and vortex briefly
5.4. Centrifuge the sample solution at 280 × g at 20°C for 1 minute
5.5. Incubate for 5 minutes at room temperature to denature the DNA into single strands.
5.6.Add 990 μlPrechilled HT1 buffer. Adding HT1 buffer results in a 20 pM denatured library in 1 mMNaOH.
5.7. Dilute the denatured DNA to the desired concentration using the following example:

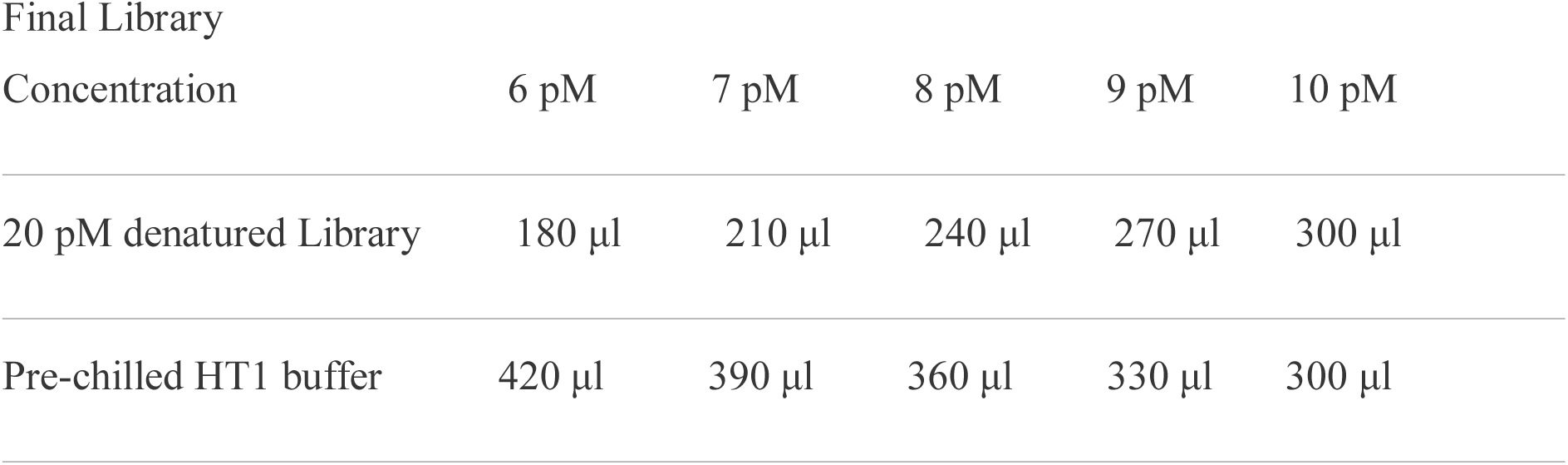

5.8. Invert several times to mix and then pulse centrifuge the DNA solution.
5.9. Place the denatured and diluted DNA on ice.
5.10. Library sequencing-Load the Denatured Library in Miseq V3 Reagent Cartridge and start the sequencing run for 2 × 300 paired end read.

### 6. Heterogeneity ‘N’ spacer Trimming

In-house python script “MetReTrim” was written to trim the heterogeneity ‘N’ spacers from the 5’ end of the reads.. Please visit the following link for details on how to download the software and other usage related information: https://github.com/Mohak91/MetReTrim. The algorithm looked for the given unique primer sequence(s) in each read and allowed upto 3 mismatches during the search. Once the primer sequence was completed, all the bases before the start of the primer sequence were trimmed. The primer sequence was retained in the processed reads. Two files were generated in the output directory-1) fastq file containing the processed reads and 2) fastq file containing unprocessed reads. The unprocessed reads were a result of primer sequences in the reads having more than 3 mismatches or insertions and deletions. The software was run as a command line using the following syntax:

For paired-end reads,

MetReTrim -i <path to fastq files folder> -o <desired path to trimmed output> -f <primer sequence for forward read> -r <primer sequence for reverse read>

## Results

The experiment was designed to obtain libraries using ‘N’ spacer primers from Metagenomic DNA and *E. coli* DNA (Fig. 1). The libraries were prepared from 5 set of N spacer primer pool combinations. The ‘N’ spacer primer pool combinations were made by equimolar pooling of forward and reverse primer (Table 3) Equimolar pool of barcoded libraries prepared using 5 set of ‘N’ spacer primer pool combinations were denatured and spiked in Hiseq 2500 Rapid-V2 2×100bp run to check base distribution at each sequencing cycle. Extracting fastq files from raw data, data de-multiplexing and Illumina adapters trimming was done using Bcl2fastq conversion software. The fastq files generated for Metagenomic DNA and *E.coli* DNA were analysed using in-house python script to check for the diversity at each base position in read 1 and read 2 sequence reads. The read 1 sequence with 6N spacer and 7N spacer (Fig. 2A, 2B and 3A and 3B) exhibited base diversity in the first ten nucleotides, allowing for better identification of clusters in the first few cycles, however at 11^th^ and 12^th^ base position, the contribution of A and C nucleotide reduced to ∼15% this lead to poor base quality scores. Also the base diversity pattern is similar for Metagenomic and *E.coli* DNA. This confirms that ‘N’ spacer primer pool is able to generate base diversity in amplicon libraries from pure culture as well. Analysis of Read1 sequence for 8N and 9N primer pool (Fig. 2C, 2D and 3C and 3D) comparatively showed more promising base diversity but nucleotide distribution at position 15^th^-16^th^ showed bias towards Green Laser Registry. Distribution of nucleotides for Green and Red laser registry plays a critical role in obtaining good quality reads, therefore Fastq results were analysed for 10N primer pool (Fig. 2E and 3E). We found that the 10N primer pool combination although showed significant increase in G nucleotide beyond 12^th^ base position the effect was balanced by elevated percentage of A and C nucleotide responsible for Red laser registry.

**Table 3:** ‘N’ spacer primer pool combination

| ‘N’ spacer primer pool combinations | Forward Primer Pool | Reverse Primer Pool |
| --- | --- | --- |
| 6 N Spacer primer pool | 6N, 5N, 4N, 3N, 2N, 1N, 0N | 6N, 5N, 4N, 3N, 2N, 1N, 0N |
| 7 N Spacer primer pool | 7N, 6N, 5N, 4N, 3N, 2N, 1N, 0N | 7N, 6N, 5N, 4N, 3N, 2N, 1N, 0N |
| 8 N Spacer primer pool | 8N, 7N, 6N, 5N, 4N, 3N, 2N, 1N, 0N | 8N, 7N, 6N, 5N, 4N, 3N, 2N, 1N, 0N |
| 9 N Spacer primer pool | 9N, 8N, 7N, 6N, 5N, 4N, 3N, 2N, 1N, 0N | 9N, 8N, 7N, 6N, 5N, 4N, 3N, 2N, 1N, 0N |
| 10 N Spacer primer pool | 10N, 8N, 7N, 6N, 5N, 4N, 3N, 2N, 1N, 0N | 10N, 8N, 7N, 6N, 5N, 4N, 3N, 2N, 1N, 0N |

**Fig. 1.**
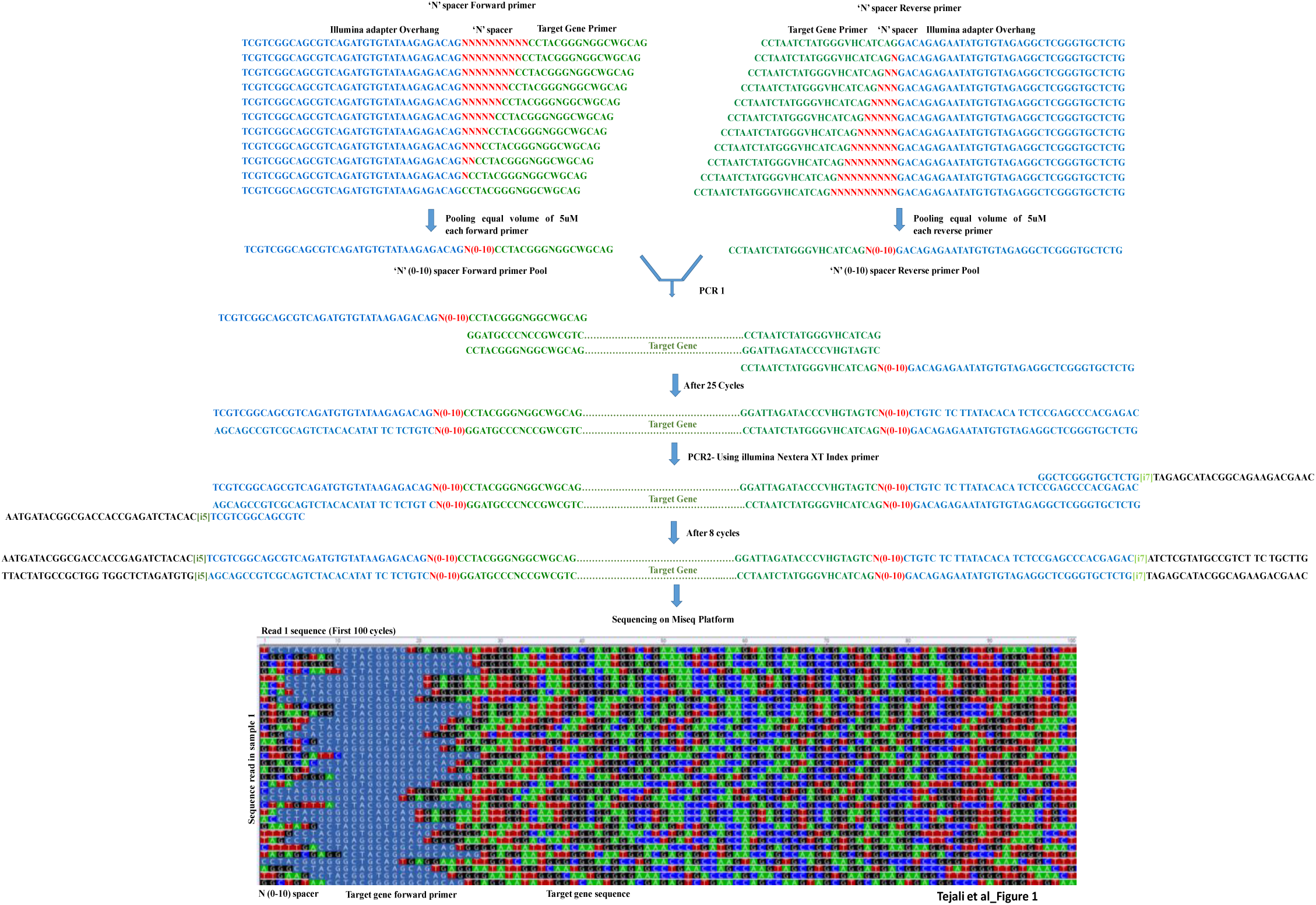
Schematic representation of 16S amplicon sequencing workflow. *E.coil* DNA was amplified for 16S V3–V4 regions using gene specific ‘N’ spacer primers with overhang adapters. The amplicons were subjected to second PCR using Nextera XT index primers. Illumina MiSeq platform was used for sequencing the final libraries. The first 100 sequencing cycles illustrates the base diversity generated within a sample amplified using ‘N’ spacer primers.

**Fig. 2.**
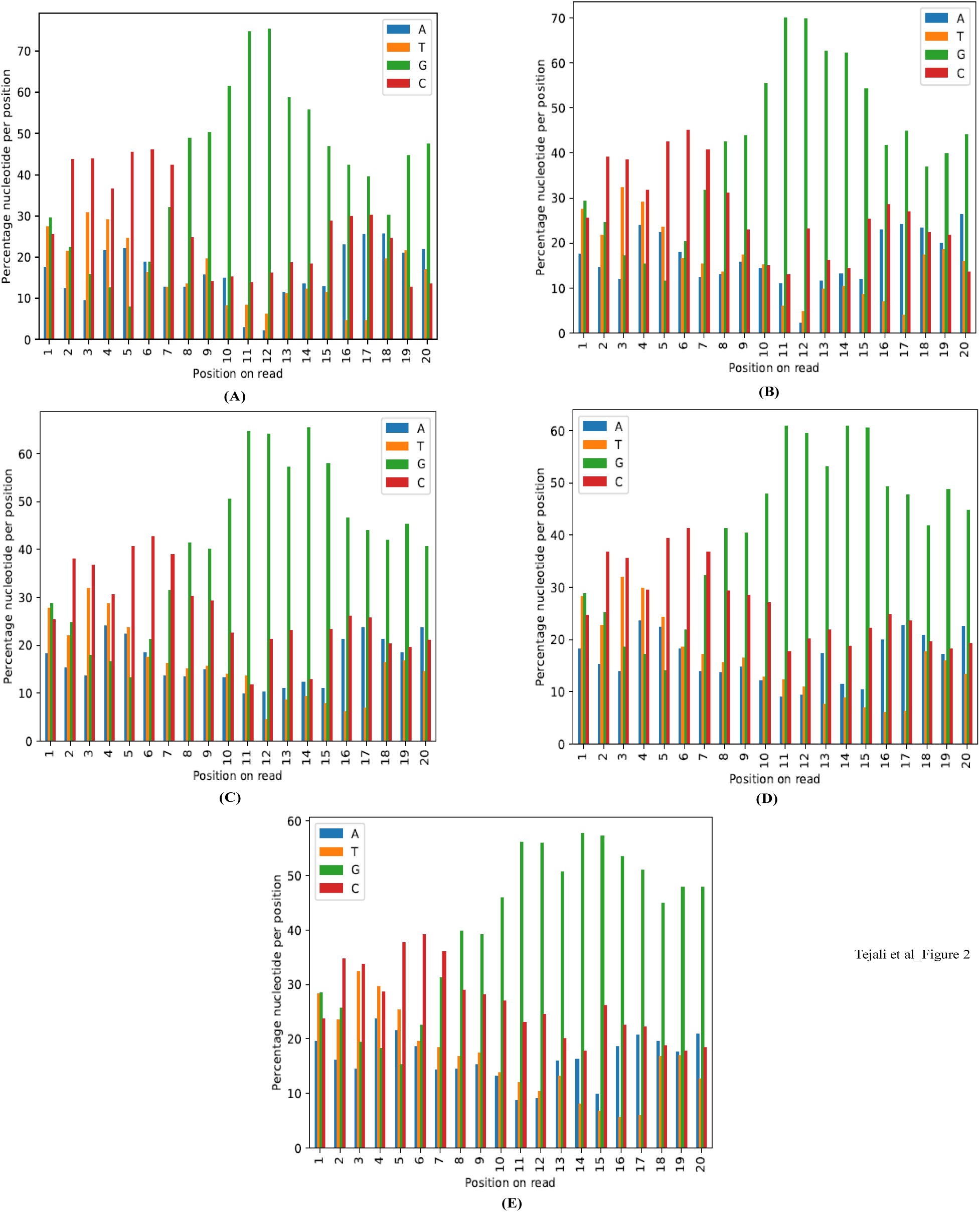
Graphical representation of base diversity in Metagenomic 16S V3-V4 amplicon sequencing using **(A)** 6N **(B)** 7N **(C)** 8N **(D)** 9N **(E)** 10N spacer primer pool sets.

**Fig. 3.**
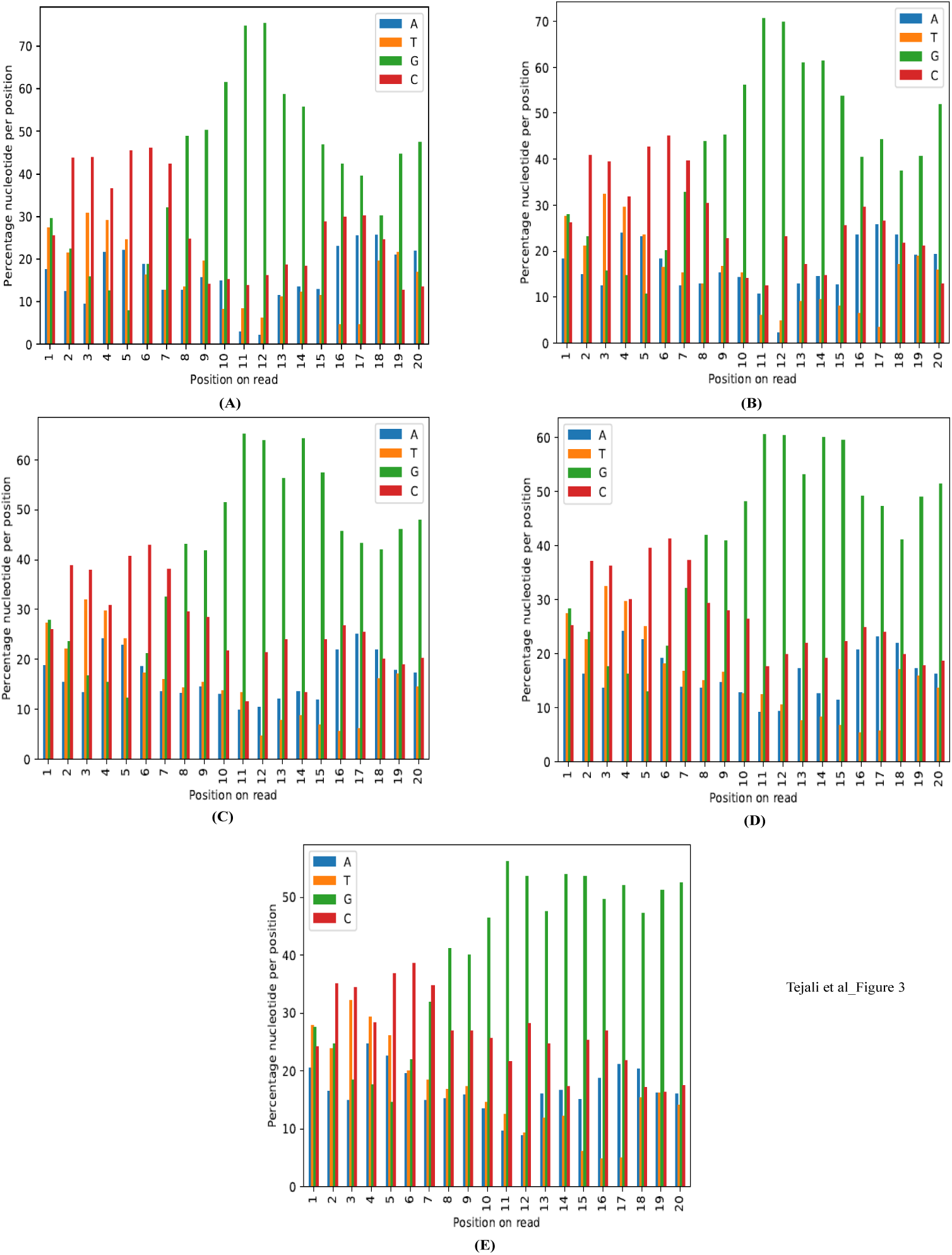
Graphical representation of base diversity in *E. coli* 16S V3-V4 amplicon sequencing using **(A)** 6N **(B)** 7N **(C)** 8N **(D)** 9N **(E)** 10N spacer primer pool sets

To evaluate the result obtained we applied our approach to prepare 16S V3-V4 amplicon library for DNA extracted from pure *E. coli* culture using 10N primer pool (Fig. 3). The library was denatured with 0.2N NaOH. Paired end sequencing was performed on illumina MiSeq platform using 2×300 V3 sequencing kit producing 290bp reads per end with loading concentration 11pM. The average quality scores (Q30) were ∼92% without PhiX spike-in. Per base sequence quality of libraries prepared using our method (‘N’ spacer V3-V4 primers) without PhiX Spike-in (Fig. 4B) showed increased sequence quality compared to libraries prepared using standard illumina V3-V4 primers with 10% PhiX spike-in (Fig 4A).

**Fig. 4.**
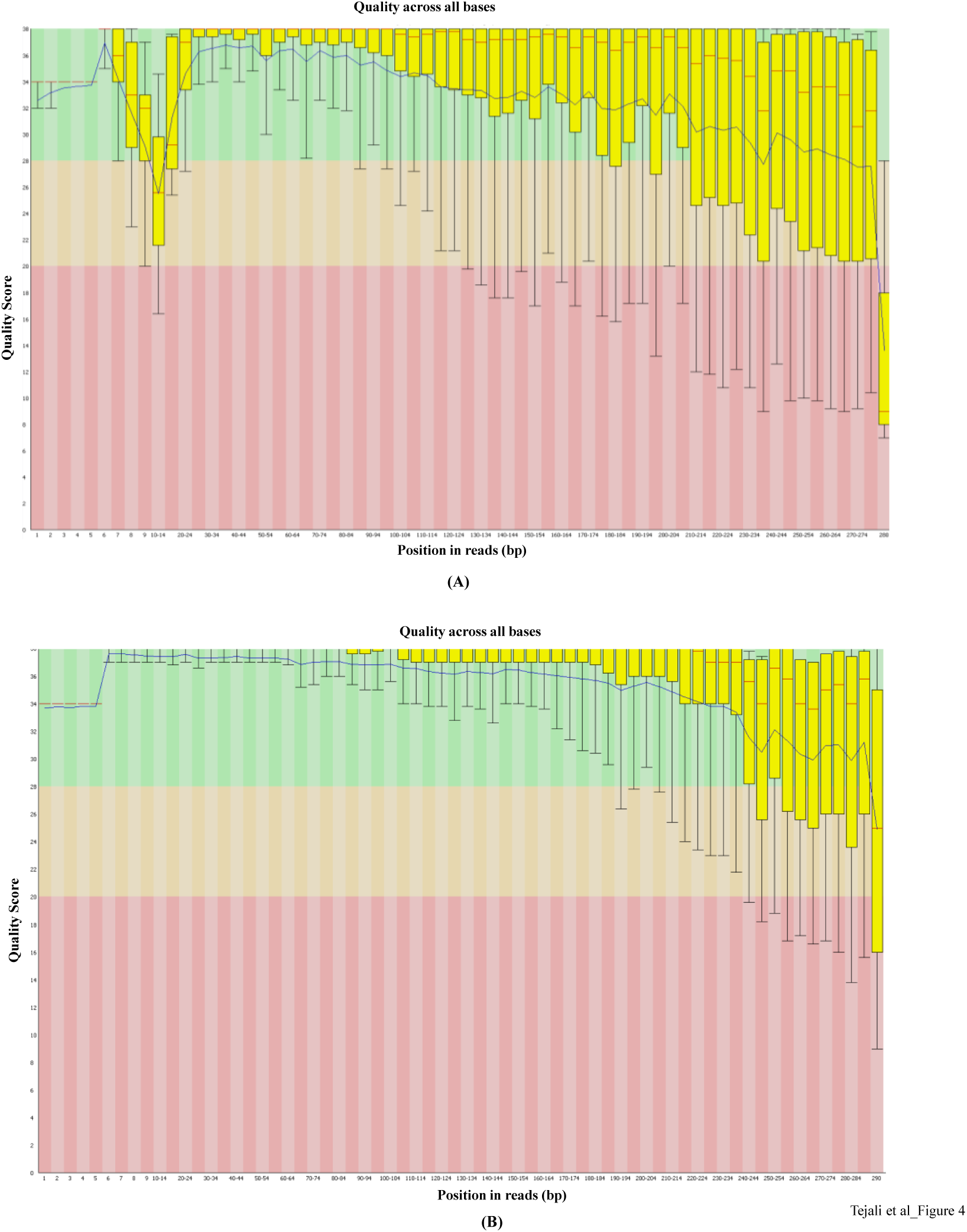
**(A)** Per base sequence quality of libraries prepared using Standard illumina V3-V4 primers with 10% PhiX spike-in. **(B)** Per base sequence quality of libraries prepared using ‘N’ spacer V3-V4 primers without PhiX Spike-in. The quality scores showed increased base sequence quality compared to libraries prepared using Standard illumina V3-V4 primers with 10% PhiX spike-in

## Comparison with the previous published amplicon sequencing protocols

Liyou et al., 2015 tried to shift the sequencing frame of amplicons by using spacers of 0–7 bases, however in our studies we observed that spacers consisting 7 bases are not sufficient to resolve the issue of unbalanced base distribution and requires higher percent of PhiX spike-in. Jensen et al., 2019 designed heterogeneity primers by adding specific nucleotide bases to 16SV3-V4 primers to create 10 oligonucleotide sets and carried out Miseq run with 10% PhiX. This approach requires preparation of minimum 10 libraries to be pooled to generate base diversity. Adopting this strategy to other amplicon library preparation needs a careful designing of heterogeneity spacers to ensure base diversity by considering the sequence of primer used to amplify the gene of interest. In our study we designed primers by adding ‘N’ spacers (0-10) and pooled them together that results in base variability within individual library leading to effective laser registry without need for PhiX spike-in. The maximum amplicon sequence length compromises for the base sacrifice by sequencing spacers. Pooling the primers for library preparation simplifies the experimental setup and designing modified primers by effortless addition of ‘N’ spacers (0-10) upstream of the target gene binding region. Jensen et al., 2019 also reported slower drop in quality scores at 185–189 position in sequence reads in contrast our method shows higher quality Q scores upto 230-240 base position and a slower drop thereafter (Fig 4B). Holm et al., 2019 presented extremely high-throughput and high-quality 300 bp paired-end reads from 1,568 amplicon libraries per lane on a HiSeq 2500 instrument with 5% PhiX spike-in. 0-7 base heterogeneity spacer strategy was used for library preparation. In our study we presented high quality paired end 2×290bp sequencing run on Miseq without PhiX spike-in and also proved that a minimum of 0-10 base heterogeneity spacer is needed to resolve the issue of base diversity.

## Conclusion

In this study, we designed heterogeneity primers by adding (0-10) ‘N’ spacers to 16S V3-V4 specific forward and reverse primer to generate amplicon with complex base diversity due to the frameshift effect. Addition of ‘N’ nucleotide to primers and pooling them together makes the experimental setup faster and simple. The use of ‘N’ spacer primers generates nucleotide distinctiveness within individual libraries at each base resulting in better identification of clusters during library sequencing run and enhance confidence in nucleotide base calls. This allows for multiplexing of more number of samples with no requirement of PhiX spike-in. This greatly reduces the overall cost and attains improved data quality. The ‘N’ spacer strategy can be used to amplify and sequence other amplicon regions. We have developed pipeline to specifically remove the heterogeneity spacers in sequenced reads.

## Acknowledgments

This work was supported by National Centre for Biological Sciences-TIFR. We thank Aswin Sai Narain Seshasayee and Dimple Notani for comments on the manuscript. We also thank NCBS, Next Generation Genomics Facility (NGGF) for their help and support.

## Author contributions

AP conceived and designed the study. TN performed all the experiments under the guidance of AP. MS performed the bioinformatics analysis. TN, MS and AP wrote the paper.

## References

[1] Lundberg Derek S., et al., Practical innovations for high-throughput amplicon sequencing. Nature methods 10.10 (2013): 999.

[2] Callahan J Benjamin et al., High-throughput ampliocn sequencing of full-length 16S rRNA gene with single-nucleotide resolution. Nucleic Acid Res 10: 47(18):2019

[3] Holm Johanna B., et al., Ultrahigh-throughput multiplexing and sequencing of> 500-base-pair amplicon regions on the IlluminaHiSeq 2500 Platform. MSystems 4.1 (2019): e00029–19.

[4] Schirmer Melanie, et al., Insight into biases and sequencing errors for amplicon sequencing with the IlluminaMiSeq platform. Nucleic acids research 43.6 (2015): e37–e37.

[5] Muinck Eric J., et al., A novel ultra high-throughput 16S rRNA gene amplicon sequencing library preparation method for the IlluminaHiSeq platform. Microbiome 5.1 (2017): 68.

[6] Illumina Low-diversity sequencing on the IlluminaHiSeq platform (Illumina Technical Note 770-2014-035). 2014

[7] Jensen Elizabeth A., et al., Heterogeneity spacers in 16S rDNA primers improve analysis of mouse gut microbiomes via greater nucleotide diversity. BioTechniques 67.2 (2019): 55–62

[8] Herbold Craig W., et al., A flexible and economical barcoding approach for highly multiplexed amplicon sequencing of diverse target genes. Frontiers in microbiology 6 (2015): 731.

[9] Kozich James J., et al., Development of a dual-index sequencing strategy and curation pipeline for analyzingamplicon sequence data on the MiSeqIllumina sequencing platform. Appl. Environ. Microbiol. 79.17 (2013): 5112–5120.

[10] Liyou, et al., Phasing amplicon sequencing on IlluminaMiseq for robust environmental microbial community analysis. BMC Microbiology 15.1 (2015): 125.

